# Overarching control of autophagy and DNA damage response by CHD6 revealed by modeling a rare human pathology

**DOI:** 10.1101/2020.01.27.921171

**Authors:** Yulia Kargapolova, Rizwan Rehimi, Hülya Kayserili, Joanna Brühl, Anne Zirkel, Yun Li, Gökhan Yigit, Alexander Hoischen, Stefan Frank, Nicole Russ, Jonathan Trautwein, Magdalena Laugsch, Eduardo Gade Gusmao, Natasa Josipovic, Janine Altmüller, Peter Nürnberg, Gernot Längst, Frank J. Kaiser, Erwan Watrin, Han Brunner, Alvaro Rada-Iglesias, Leo Kurian, Bernd Wollnik, Karim Bouazoune, Argyris Papantonis

## Abstract

Members of the chromodomain-helicase-DNA binding (CHD) protein family are chromatin remodelers critically implicated in human pathologies, with CHD6 being one of its least studied members. Here, we discovered a *de novo CHD6* missense mutation in a patient clinically presenting the rare Hallermann-Streiff syndrome (HSS). We used genome editing to generate isogenic iPSC lines and model HSS in relevant cell types. We show that CHD6 binds a cohort of autophagy and stress response genes across cell types. The HSS-mutation affects CHD6 protein folding and impairs its ability to recruit co-factors in response to DNA damage or autophagy stimulation. This leads to an accumulation of DNA damage burden and to senescence-like phenotypes. By combining genomics and functional assays, we describe for the first time a molecular mechanism for the chromatin control of autophagic flux and genotoxic stress surveillance that applies broadly to human cell types and explains HSS onset.

## INTRODUCTION

Modulation of DNA accessibility is central to the regulation of eukaryotic genome functions like gene transcription, DNA replication or repair^1,2,3^. One large class of enzymes, the ATP-dependent chromatin remodeling factors, can alter DNA accessibility by removing or repositioning nucleosomal proteins along chromosomes. In mammals, the chromodomain-helicase-DNA binding (CHD) proteins represent the largest family of remodelers. In humans, all nine of its members are characterized by the presence of tandem N-terminal chromodomains and by a central SNF2-like ATPase module. Additional domains allow for the further classification of these large (>200 kDa) proteins into three subfamilies. Subfamily I includes CHD1 and −2 that carry DNA-binding domains that are absent from subfamily II members CHD3-5; these instead have tandem plant homeodomains to recognize and bind histone tails. Subfamily III includes CHD6-9, marked by the presence of Brahma/Kismet (BRK) and SANT-like domains closer to their C-termini^4,5^. SANT domains, initially found in co-repressor proteins, were later also identified in different remodeling complex subunits as modules allowing binding to histones concomitantly with enzymatic catalysis^6^. The CHD SANT/SLIDE domains, which resemble Myb DNA-binding domains, are conserved between CHDs and ISWI proteins. Data suggest that the SANT and SLIDE modules interact with DNA as one cooperative unit important for tuning DNA binding and nucleosome spacing^7^.

Mutations in subfamily III members have been causally implicated in autism (CHD7 and −8)^8,9,10^ and in the CHARGE syndrome (CHD7)^11,12,13^. CHD6 is a far less studied member of this subfamily. It is ubiquitously-expressed and found alongside RNA polymerases at nucleoplasmic sites of nascent RNA synthesis as part of supramolecular complexes^14^. Recent reports have implicated CHD6 in the repression of viral replication^15^, in the topological organization of the *CFTR* locus^16^, and in chromatin remodeling at sites of oxidative DNA damage^17^. To date, CHD6 has not been functionally linked to any human pathology, but there exist reports of large translocations that also include its locus in one Pitt-Hopkins patient^18^, in a single case of mental retardation^19^, and in sporadic acute myeloid leukemia incidences^20^.

Throughout development, human cells continuously face genotoxic stress and mechanisms are in place to survey and restore the resulting DNA damage. Weakening of these mechanisms leads to DNA damage accumulation and is now understood to cause premature ageing syndromes (known as segmental progerias)^22^. For example, the well-studied Hutchinson-Gilford and Werner progerias stem from mutations in *LMNA* and *RECQL2*, respectively, which in turn promote genome instability^23,24^. Alongside increased DNA damage burden, decreased autophagy levels constitute another hallmark of aging. Autophagy is a housekeeping catabolic pathway essential for recycling long-lived organelles and misfolded proteins, and is activated by various stimuli, including growth factor withdrawal, nutrient deprivation, infection, oxidative stress or hypoxia^25^. A functional link between autophagy induction and DNA damage recognition and repair was recently documented^26,27^. Interestingly, mouse models with conditional, tissue-specific knockout of key autophagy regulators present age-associated defects^28^.

The Hallermann-Streiff syndrome (HSS; OMIM ID: #234100) is a rare congenital disorder characterized by craniofacial and dental dysmorphisms with a specific facial gestalt, eye malformations, distinctive facial features, hair and skin abnormalities, and short stature. Due to its clinical course and progression, HSS is regarded as a premature aging disorder^29,30,31^. With only few cases reported to date, and with virtually all reports being descriptive, there is an apparent need for dissecting the molecular pathways underlying HSS^32^. Given that in the only available mouse model for *CHD6* lacking exon 12 (encoding its conserved ATPase domain) no obvious phenotype apart from mild ataxia is observed^21^, we chose to model this disease by generating isogenic induced pluripotent stem cell (iPSC) lines carrying or not the *CHD6* mutation. Using these lines in genomics and functional studies, we provide the first molecular insights into HSS etiology. We identify CHD6 as a major and ubiquitous upstream regulator of stress and autophagy response genes across cell types. The HSS mutation interferes with co-factor recruitment to CHD6 target genes, resulting in impaired autophagy flux, DNA damage accumulation, and development of senescence-like hallmarks.

## RESULTS

### A putatively-causative HSS mutation and generation of isogenic iPSC lines

In order to understand the molecular basis of HSS, we sought to identify genetic mutations associated with the disease. Despite the rarity of HSS samples, whole exome sequencing of blood and saliva-derived DNA of a patient and parents uncovered a single *de novo* missense mutation resulting in an isoleucine to methionine (I1600M) amino acid exchange in the *CHD6* coding sequence. This putatively-causative heterozygous mutation in *CHD6* maps to its predicted second SLIDE domain, at a position highly conserved across species and also present in the equivalent SLIDE domain of CHD1 (**Fig. 1a** and **S1a**). We derived iPSCs from this HSS patient by reprograming fibroblasts from a skin biopsy. Successful reprograming was confirmed using pluripotency markers at the RNA and protein levels (**Figs 1b** and **S1b**). We confirmed the ability of patient-derived iPSCs to form embryonic bodies (**Fig. S1c**) and to spontaneously differentiate into all three germ layers (**Fig. S1d-f**). As controls for these tests, we used previously characterized age- and sex-matched wild-type iPSCs^33^. However, these lines do not have identical genomic backgrounds, which might complicate data interpretation. To address this, we used different CRISPR-Cas9 editing approaches on our control iPSCs to generate isogenic lines carrying the *CHD6* mutation. We obtained two independent lines expressing only the mutated *CHD6* allele (hereafter “mut”), and another two lines heterozygously expressing wild-type and mutated *CHD6*, like in the HSS patient (hereafter “het”; **Figs 1c** and **S1g**). These lines were diploid and karyotypically stable, and could be differentiated into neural crest cells (NCCs) and spontaneously-contracting cardiomyocytes (CMs) with comparable efficiencies (**Figs 1d,e** and **S1h,i**). NCCs and CMs were chosen due to their relevance to this syndrome, since progressing cardiomyopathy is a recurrent HSS manifestation and it also invariably presents as a neurocristopathy^32^. Taken together, we now have a model to study the mechanisms underlying HSS, while also characterizing CHD6 in both a proliferative and a post-mitotic context.

**Fig. 1.**
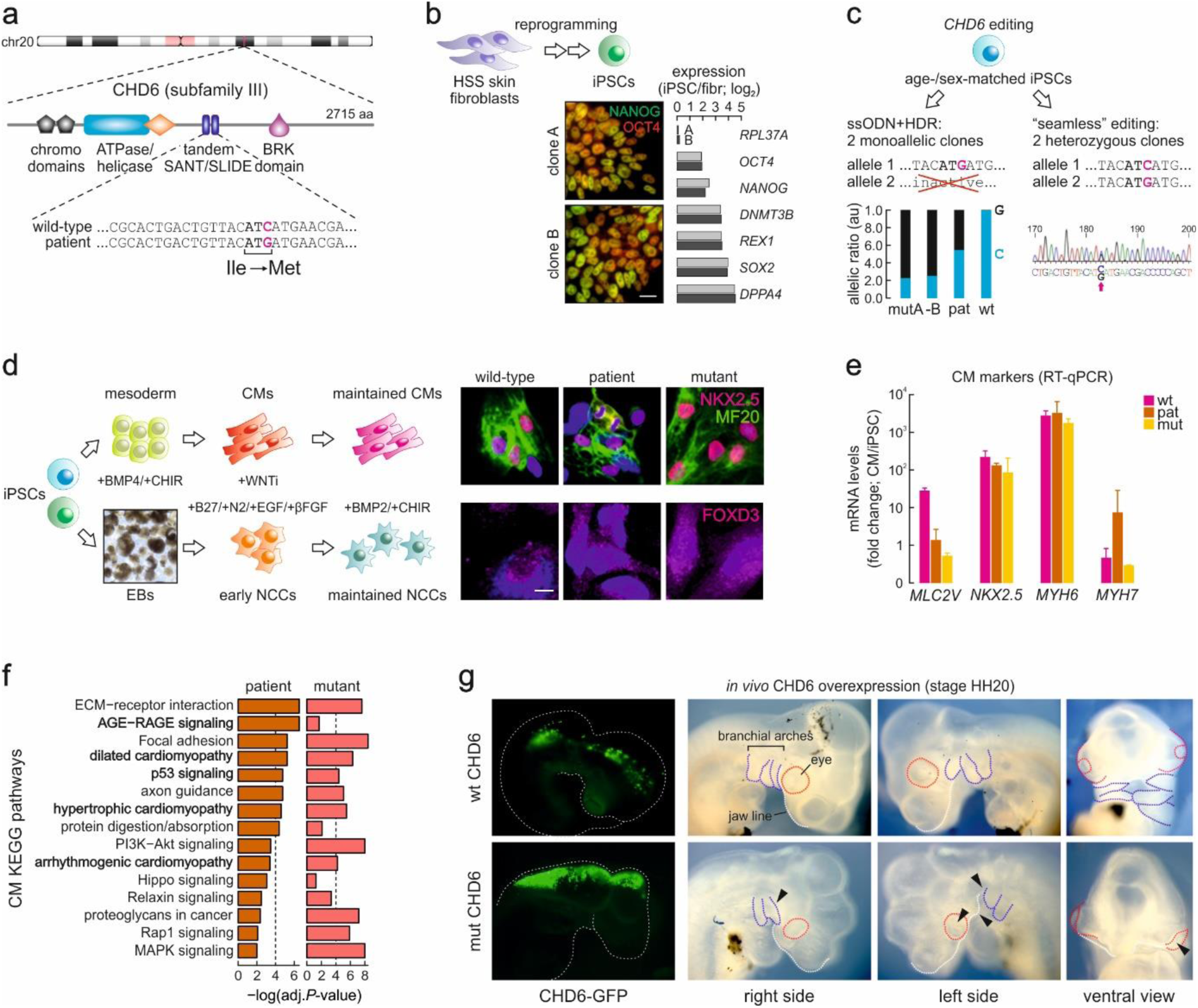
Generation of isogenic iPSC lines expressing the I1600M CHD6 mutant. (**a**) Schematic representation of the *CHD6* locus (*top*) highlighting key functional protein domains (*middle*) and the HSS-specific I1600M missense mutation in the second SANT/SLIDE domain (*bottom*). (**b**) Nuclear reprogramming of skin fibroblasts from an HSS patient carrying the I1600M *CHD6* mutation. iPSC identity is exemplified by immunofluorescence (*bottom left*) or RT-qPCR of pluripotency markers (*bottom right*) in two independent clones. (**c**) Generation of iPSCs monoallelically (*left*) or heterozygously-expressing I1600M CHD6 (*right*) using different CRISPR/Cas9 genome editing strategies. (**d**) Derivation of wild-type (wt), homozygous mutant (mut), or patient-derived (pat) cardiomyocytes (CMs; *top row*) or neural crest cells (NCCs; *bottom row*) from iPSCs exemplified by immunofluorescence detection of lineage-specific marker genes (*right*). (**e**) RT-qPCR data showing mean CM marker gene expression levels (±SD; relative to iPSC levels) from all genotypes in panel d. (**f**) Top 15 KEGG pathways commonly misregulated in patient (*left*) and mutant CMs (*right*) according to RNA-seq data. (**g**) *In vivo* overexpression of wild-type (*top row*) and mutant CHD6 (*bottom row*) in chicken embryos at stage HH20 and examination of lateral and ventral views for the development of branchial arches (*blue*), eyes (*red*), and jaw (*white*).

### The I1600M CHD6 mutation affects gene expression and development

To understand how the *CHD6* mutation affects gene expression, we generated transcriptome profiles using total cell RNA-seq from all genotypes in both NCCs and CMs. Gene set enrichment analysis against the KEGG pathway database revealed that both patient-derived and gene-edited mutant CMs were affected as regards ECM interactions, focal adhesion and AGE-RAGE/p53 signaling. It also highlighted that genes involved in dilated, arrhythmogenic or hypertrophic cardiomyopathy were deregulated (**Fig. 1f**). Looking for genes commonly deregulated between patient-derived CMs and NCCs, we found those associated with axon guidance, lipid biosynthesis, heart contraction to be consistently downregulated compared to wild-type levels (**Fig. S1k**). On the other hand, genes linked to cell adhesion, wound healing, apoptotic signaling, and the integrin pathway were found upregulated (**Fig. S1j**).

Given that focal adhesion and integrin components are important for proper cell differentiation and specification, we hypothesized that, especially for NCCs, the CHD6 mutation could lead to altered migratory patterns. Although we did not observe significant differences in *in vitro* migration assays (**Fig. S1l**), we also tested this hypothesis in chicken embryos, an established model for NCC studies. We overexpressed human wt or HSS-mutant CHD6 directly in the chicken neural tube via electroporation of relevant constructs. We examined the resulting embryo phenotypes at developmental stage HH20, where wtCHD6-expressing cells displayed the expected migration to the cornea, heart, and along the neural tube (**Fig. 1g**, *top row*). In contrast, mutCHD6-expressing cells accumulated more in the neural tube and brain of the embryo. Moreover, these embryos displayed obvious facial malformations, lack of the 1^st^ or fusion of the 1^st^ and 2^nd^ branchial arches, enlarged forebrain and heart, as well as diminished mesenchymal tissue, all of which are reminiscent of HSS phenotypes (**Fig. 1g**, *bottom row*). As a whole, our data demonstrate how a single *CHD6* mutation critically impacts development and gene expression.

### CHD6 binds autophagy-relevant genes across cell types

The binding preferences of CHD6 along human chromosomes remain unknown. To address this and infer CHD6-regulated loci, we applied a tailored crosslinking and ChIP protocol (see **Methods**) to iPSCs, NCCs, and CMs of all genotypes (**Figs 2a** and **S2a**). This ChIP-seq data collection revealed two unexpected features. First, mutCHD6, irrespective of the cell type studied, consistently displayed stronger peaks and a broader binding catalogue than wtCHD6 (**Figs 2a-c** and **S2a-c**). Second, CHD6 binds many of the same loci across three diverse cell types, mostly at gene promoters (**Figs 2c,d** and **S2a**). CHD6-bound loci are associated with autophagy, cell cycle regulation, and the DNA damage response (**Figs 2e** and **S2d**). While CHD6 partially overlaps CHD2-bound positions, it does not generally coincide with CHD1 or CHD7, indicating it regulates disparate pathways (**Fig. S2e**).

**Fig. 2.**
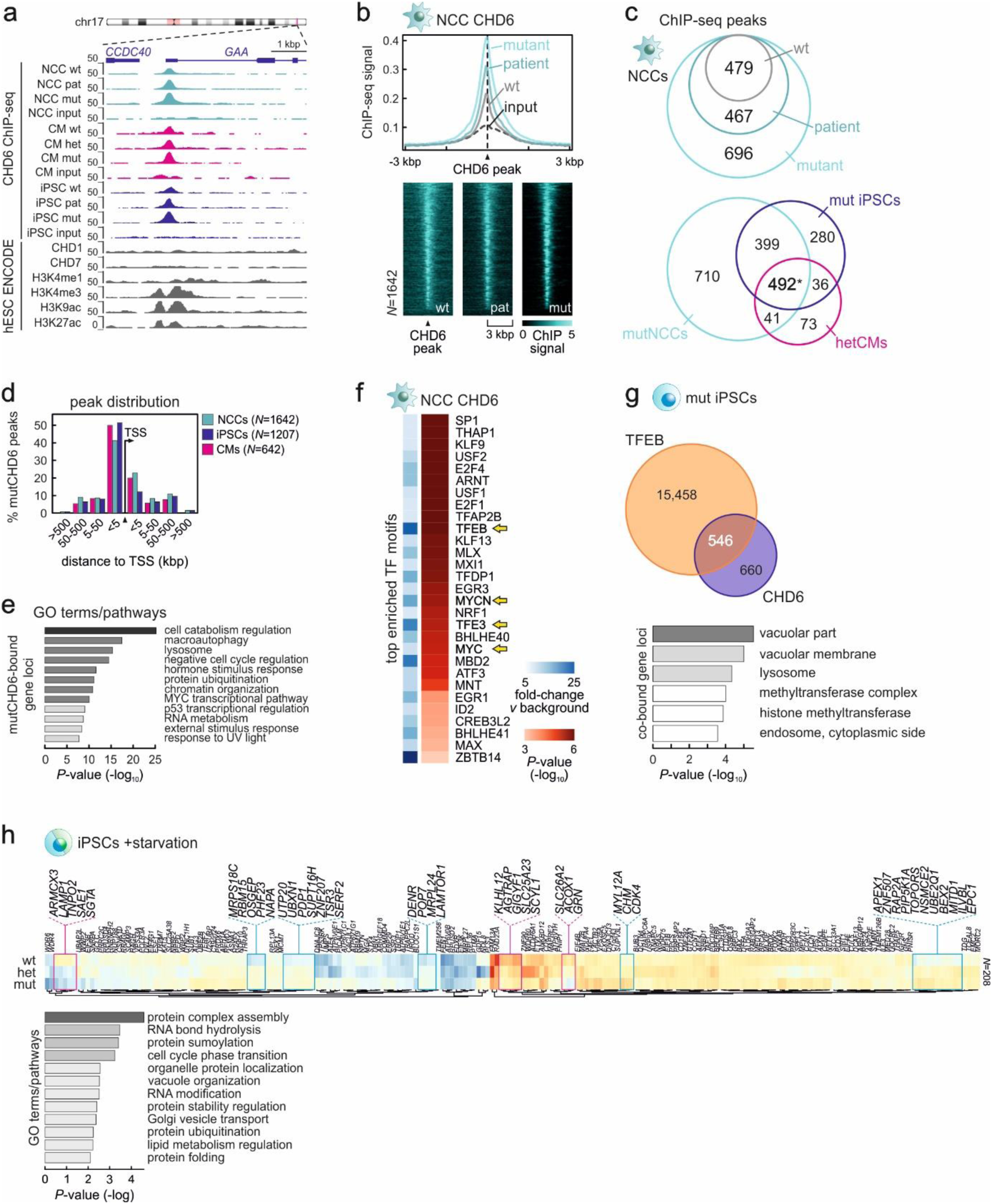
CHD6 binding properties along human chromosomes. (**a**) Genome browser views of CHD6 ChIP-seq data from wild-type (wt), monoallelic mutant (mut), heterozygous mutant (het) or patient-derived (pat) NCCs (*green*), CMs (*magenta*), and iPSCs (*blue*) around the *GAA* promoter. Reference hESC ENCODE ChIP-seq data are also aligned below. (**b**) Line plot (*top*) and heatmaps (*bottom*) showing ChIP-seq signal distribution in the 6 kbp around CHD6-bound sites from wild-type (*grey*), patient-derived (*dark green*), or homozygous mutant NCCs (*light green*). (**c**) Venn diagrams showing overlap of CHD6 ChIP-seq peaks from panel b (*top*) or from three cell types (*bottom*). \**P*<0.0001; significantly more than expected by chance according to a hypergeometric test. (**d**) Bar plot showing the percentage of mutCHD6 ChIP-seq peaks located at increasing distances up- or downstream of gene TSSs across all three cell types. (**e**) Bar plot showing significantly enriched GO terms associated with mutCHD6-bound genes in NCCs. (**f**) Heatmaps showing enrichment of TF motifs (over background; *light to dark blue*) and associated *P*-values (*light to dark red*) within accessible DNase I footprints under CHD6 ChIP-seq peaks. (**g**) Venn diagram showing overlapping TFEB and CHD6 peaks from mutant iPSCs (*top*) and the top most significantly-enriched GO terms associated with the genes under the 546 shared peaks. (**h**) Heatmaps (*top*) showing changes in mRNA levels (log_2_) of genes differentially regulated in wild-type iPSCs upon 2-h starvation. Those convergently up-/downregulated in monoallelic- (mut) and heterozygous-mutant (het) cells are highlighted (*magenta/blue rectangles*). Bar plot (*bottom*) showing significantly-enriched GO terms associated with the highlighted genes.

Analysis of transcription factor binding motifs in accessible DNase I footprints under CHD6 peaks returned TFEB and TFE3 recognition sequences as two of the most enriched (**Fig. 2f**). In addition, survey of ENCODE ChIP-seq data found TFEB as significantly enriched TF at CHD6-bound sites (**Fig. S2f**). TFEB and TFE3 are major regulators of autophagy and lysosomal genes^34^. Accordingly, we performed TFEB ChIP-seq in iPSCs and confirmed it overlaps ~45% of CHD6 peaks (**Figs 2g** and **S2g**). Genes bound by both TFEB and CHD6 were associated with GO terms like lysosome, ER stress response, and cell catabolism (all processes linked to autophagy; **Fig. 2g**). Thus, mutCHD6 could affect expression profiles in response to pro-autophagy cues. To test this, we subjected wt-, het- and mut-iPSCs to 2-h starvation and performed 3’-end RNA-seq. Wild-type iPSCs responded via downregulation of mitochondrial and ribosomal processes, while mutant cells did not. Mut/het-iPSCs also activated proinflammatory genes, ribosome biogenesis and suppressed DNA damage response ones (**Fig. S2h**). Moreover, ERK1/2 signaling that is known to regulate TFEB subcellular localization^35^ was also affected. Looking specifically at CHD6-bound genes, >200 showed levels deviating from wild-type in response to starvation (**Fig. 2h**, *top*). Of these, the genes that showed similar misexpression trends in both het- and mut-iPSCs were linked to processes like RNA hydrolysis and modification, cell cycle transition, vacuole organization, and protein turnover (**Fig. 2h**, *bottom*). Together, our data assign a key role to CHD6 in regulating autophagy genes.

### CHD6 mutation leads to impaired autophagy and increased DNA damage burden

To assess the functional involvement of CHD6 in autophagy regulation, we stained cells for some of its key regulators. First, localization and quantification of microtubule-associated LC3 showed it is depleted from CM nuclei only in the context of the *CHD6* mutation (**Fig. 3a**). This should not occur in the absence of pro-autophagy cues as nutrient deprivation is what drives LC3 deacetylation by SIRT1 and cytoplasmic translocation. However, we documented elevated SIRT1 levels in both patient-derived and het-CMs (**Fig. 3b**). Likewise, the decreased levels of phosphorylated ribosomal S6 proteins in mutant CMs suggest dampened mTORC1-pathway activity and autophagy induction (**Fig. 3c**). Critically though, these marks of activated autophagy flux are counteracted by significantly reduced levels of the key autophagy regulator p62 (**Fig. 3d**). Accordingly, iPSCs with mutant *CHD6* alleles show accumulation of ATG12/ATG5 dimers that are necessary for autophagy activation^34^, but have significantly fewer lysosomes (**Fig. S3a,b**). Together, our data confirm a hindered autophagy flux and turnover. Interestingly, NCCs show similar trends as regards LC3 levels, but elevated phospho-S6 (**Fig. S3c,d**). This possibly signifies intensified glycolysis via partial mTORC1 activation on top of autophagy impairment, highlighting differences between the proliferating and post-mitotic contexts.

**Fig. 3.**
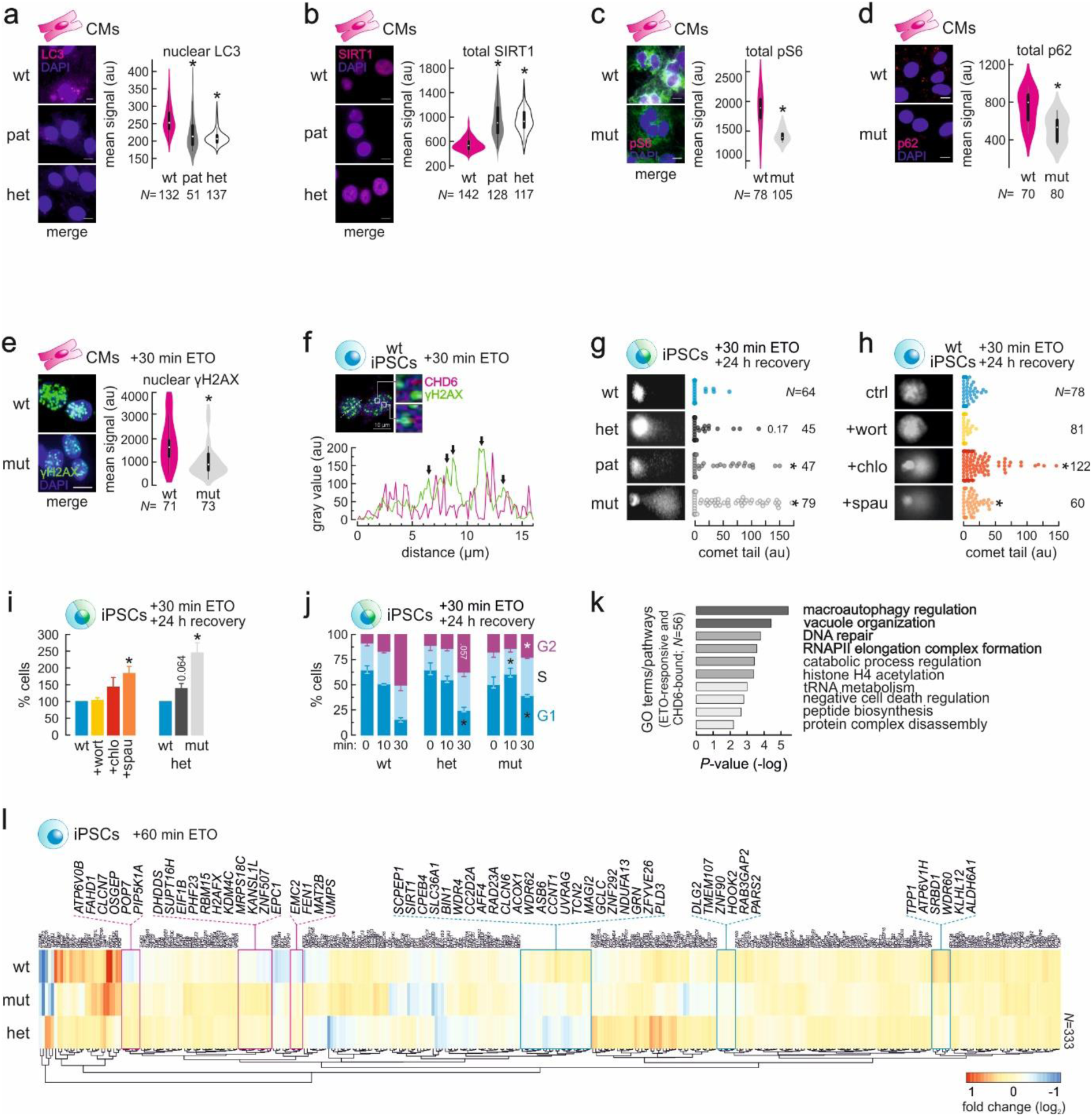
CHD6 controls autophagy and affects the DNA damage response. (**a**) Representative immunofluorescence images (*left*) of wild-type (wt), monoallelic mutant (mut), heterozygous mutant (het) or patient-derived (pat) CMs stained for LC3 (*magenta*), and bean plots quantifying nuclear signal (*right*). The number of cells analyzed (*N*) are given below each plot. *: significantly different to wild-type; *P*<0.05, Wilcoxon-Mann-Whitney test. (**b**) As in panel a, but for total SIRT1 levels in CMs. (**c**) As in panel a, but for total phospho-S6 levels in CMs. (**d**) As in panel a, but for total p62 levels in CMs. (**e**) As in panel a, but for nuclear γH2A.X levels in CMs. (**f**) Representative immunofluorescence image of wild-type iPSCs stained for γH2A.X (*green*) and CHD6 (*magenta*) after 30-min of etoposide treatment (*top*), and line scan of signal profiles (*bottom*); γH2A.X signal foci are indicated (*arrows*). (**g**) Comet assays and quantification of tails in wild-type (wt), monoallelic mutant (mut), heterozygous mutant (het) or patient-derived (pat) iPSCs treated with etoposide for 30 min and allowed 24 h to recover; numbers of cells analyzed (*N*) are given. *: significantly different to wild-type; *P*<0.05, Wilcoxon-Mann-Whitney test. (**h**) As in panel g, but for wild-type iPSCs treated with etoposide and allowed to recover for 24 h in the presence or absence of three autophagy inhibitors. (**i**) Bar plots showing the percentage (±SD) of wild-type (wt), monoallelic mutant (mut), heterozygous mutant (het) or patient-derived (pat) that survived apoptosis, also in the presence or absence of autophagy inhibitors. *: significantly different mean compared to wild-type; *P*<0.05, two-tailed unpaired Student’s t-test. (**j**) Bar plots showing the percentage (±SEM) in the G1, S or G2 cell cycle phase upon 0, 10 or 30 min of etoposide treatment and 24 h recovery in wild-type (wt), monoallelic mutant (mut) or heterozygous mutant (het) isogenic iPSCs; *: significantly different to wild-type; *P*<0.05, two-tailed unpaired Student’s t-test. (**k**) Bar plot showing significantly-enriched GO terms associated with the genes highlighted in panel l. (**l**) Heatmap showing changes in mRNA levels (log_2_) of genes that are differentially regulated in wild-type iPSCs upon 1h etoposide treatment. Those convergently up-/downregulated in monoallelic (mut) and heterozygous mutant (het) cells are highlighted (*magenta/blue rectangles*).

Previous studies have shown that autophagy is essential for implementing DNA damage responses^28,26,27^. Thus, we examined whether impaired autophagy flux seen in *CHD6-*mutant cells affects DNA damage response and repair. We used etoposide to induce DNA double strand breaks in CMs and observed reduced Ser139-phosphorylated γH2A.X levels and suboptimal formation of γH2A.X foci in mutant cells (**Fig. 3e**). Interestingly, CHD6 does not redistribute in the nucleus of wild-type cells in response to etoposide treatment, nor does it colocalize with γH2A.X foci (**Fig. 3f**). Using iPSC extracts in western blots (given the ubiquitous role of CHD6, iPSCs are used here for convenience), we observed markedly reduced activation (via phosphorylation) of several components of the DNA damage response pathway, like p53, CHK1/2 and γH2A.X in *CHD6-*mutant cells (**Fig. S3a**). In comet assays, pronounced accumulation of unrepaired DNA was seen in iPSCs carrying mutant *CHD6* alleles, with cells expressing no wtCHD6 being most susceptible (**Fig. 3g**). Such accumulation of unrepaired breaks was recapitulated in wt-iPSCs when etoposide treatment was combined with pharmacological inhibition of autophagy (**Fig. 3h**). Notably, we obtained such results when inhibiting late autophagy stages (autolysosome formation and degradation) via chloroquine or spautin, but not when using wortmanin (targeting autophagosome maturation; **Fig. 3h**).

This impaired response to DNA damage due to the HSS-specific *CHD6* mutation is also reflected in various cellular functions. First, mutant iPSCs are significantly less susceptible to apoptosis, and this prosurvival effect was recapitulated in part upon autophagy inhibition in wt-iPSCs (**Fig. 3i**). Second, cell cycle analysis showed that mut-iPSCs are insensitive to G2/M-phase arrest despite increased DNA damage burden (**Fig. 3j**), most likely due to impaired signaling (**Fig. S3a**). Third, gene expression following DNA damage induction is also affected, as revealed by 3’-end RNA-seq on our isogenic iPSC lines. CHD6-bound genes deregulated in both *CHD6*-mutant backgrounds were especially enriched for regulators of macroautophagy, vacuole and cell catabolism regulation, RNAPII elongation, DNA repair, and cell death (**Fig. 3k,l**). Finally, we could detect hallmarks of a senescence-like phenotype in cells carrying mutant *CHD6* alleles, most likely due to damage accumulation^36,37^. The presence of mutCHD6 coincided with reduced levels of H3K27me3-marked heterochromatin (**Fig. S3e**). Similarly, mut-/het-iPSCs displayed an emergence of HP1α foci in constitutive heterochromatin, as well as strong β-galactosidase staining, both recapitulated via autophagy inhibition using spautin in wt cells (**Fig. S3f,g**). Collectively, we show that CHD6 is needed for the proper control of autophagy-related genes, and consequently for efficient DNA damage response.

### CHD6 in vitro properties are not affected by the HSS mutation

CHD6 is a chromatin remodeling factor and the HSS-relevant I1600M mutation maps inside its CHDCT2 domain (**Fig. S1a**). This domain was initially identified in CHD4 and then in subfamily II members CHD3 and −5^38^. Low stringency query of the NCBI Conserved Domain Database (max. E-value=100) uncovered that the CHD6 CHDCT2 is in fact related to the SLIDE (SANT-Like ISWI) domain implicated in DNA binding ^6,39,7^. We therefore tested whether the I1600M mutation affected CHD6 binding *in vitro* using purified full-length wt and mutant CHD6, expressed via a baculovirus system (**Fig. S4a**). Electrophoretic mobility shift assays (EMSA) show that binding to a nucleosome or to DNA (**Fig. S4b,c**) by mutCHD6 was virtually identical to wild-type. This is consistent with both our ChIP-seq data (see **Figs 2a,b** and **S2a,b**), and with the fact that the HSS mutation does not impact a lysine or arginine residue, shown to be important for DNA binding by SANT-SLIDE modules^7^. To exclude that additional DNA binding modules in CHD6 like its chromodomains^40,41^ might compensate for the I1600M mutation, we purified and tested the second CHD6 SANT-SLIDE module alone in EMSAs. Again, wt and mutant CHD6 SANT-SLIDE modules did bind chromatin with comparable efficiencies (**Fig. S4d**).

We next tested whether the I1600M mutation impinged on CHD6 remodeling activity. Using restriction enzyme accessibility (REA) assays, we observed that the ability of mutCHD6 for exposing RE sites was indistinguishable from that of the wt protein. This held true regardless of whether the RE site was near the entry/exit nucleosome site (*Mfe*I site at +28 bp) or near the pseudo-dyad axis (*Hin*6I at +71 bp; **Fig. S4e**). The ability of CHD6 to expose *Hin*6I sites should require extensive unwrapping or sliding of the nucleosome. This contrasts recent work showing that CHD6, unlike all other subfamily III members, did not slide nucleosomes *in vitro*, but only disrupted DNA-histone association in a non-sliding manner^42^. Thus, we revisited this aspect using sliding assays. Addition of full-length wtCHD6 to a nucleosome positioned at the end of a 227-bp DNA fragment resulted in the emergence of discrete slower-migrating bands, indicative of histone octamer repositioning in an ATP-dependent manner (**Fig. S4f**). Consistent with REA assays, the activity of mutCHD6 was indistinguishable from that of the wild-type protein. Finally, as CHD6 mostly binds to TSSs between the −1 and +1 nucleosomes (**Fig. S4g**), we selected a set of CHD6-bound loci to assess changes in DNA accessibility in iPSCs. We designed primers overlapping nucleosomes directly downstream of CHD6 peaks and used them in MNase-qPCR assays on mononucleosomal templates (**Fig. S4h,i**). All positions analyzed showed weak (~20%) to significant (~40%) changes in DNA accessibility (**Fig. S4h**), in line with differences inferred from our ChIP-seq data. In summary, we show that CHD6 contributes in setting up chromatin configuration at its target loci.

### Mutated CHD6 fails to recruit co-factors and activate autophagy genes

The above data suggest that the I1600M mutation in CHD6 does not impede its association to chromatin or nucleosome sliding. However, strong regulatory effects were seen across all of our HSS-mutant lines. To reconcile these, we first used structure prediction tools, namely I-TASSER^43^, HHpred^44^ and RaptorX^45^, and uncovered that the CHD6 region between aa 1448-1608 can fold into a structure highly similar to that of the Chd1 SANT-SLIDE domain^41^ (**Fig. 4a**, *left*). Introducing the HSS mutation in this *in silico* test returned a distorted interface on the modeled domain (**Fig. 4a**, *right*). In line with this data, nanoDSF measurements of full-length CHD6 showed that the mutant unfolds at significantly lower temperatures (mean T*m*: ~41.5°C) compared to the wild-type protein (~47.7°C; **Fig. 4b**). Taken together, our data strongly suggest that the I1600M mutation leads to reorganized CHD6 folding.

**Fig. 4.**
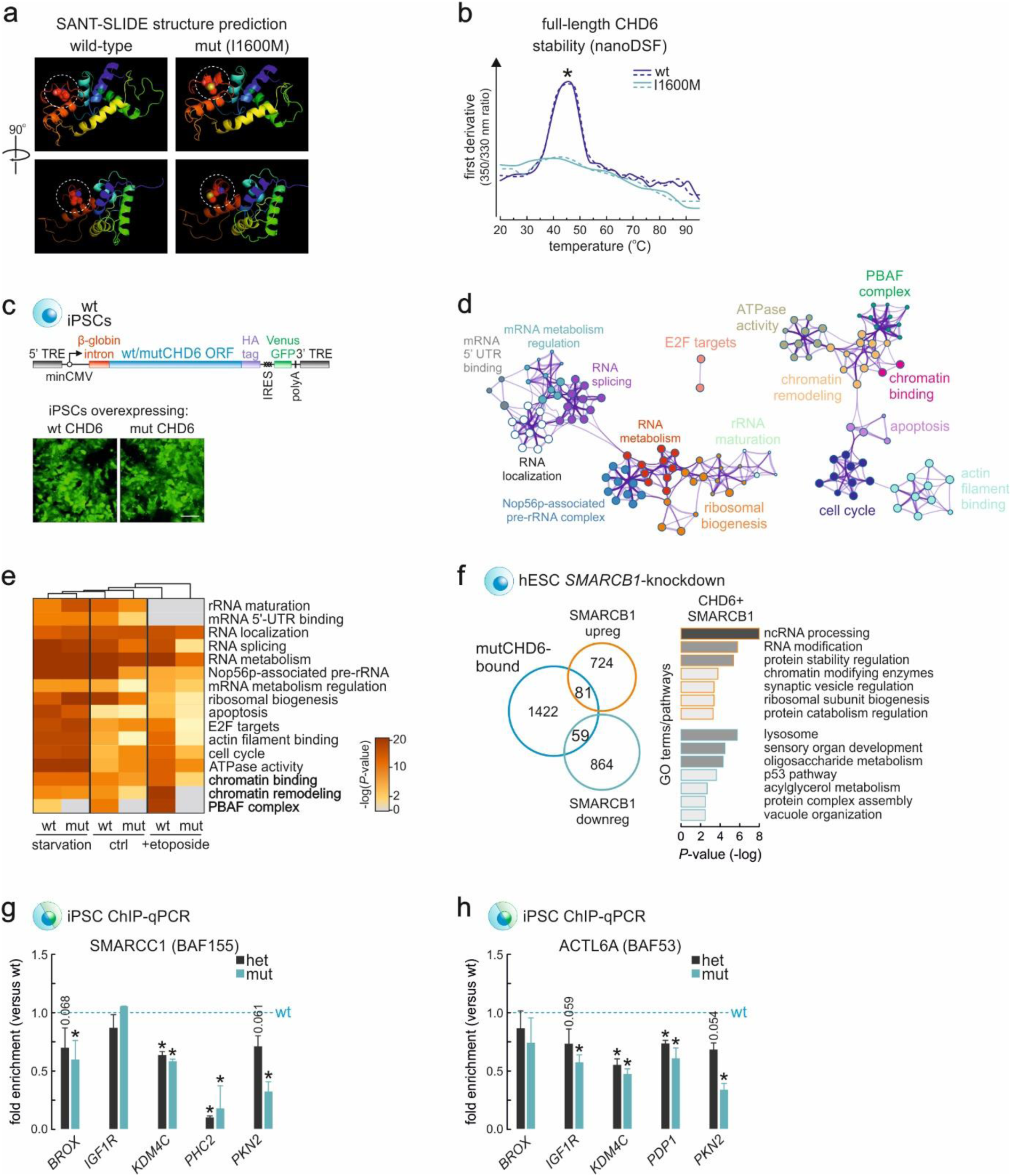
Mutant CHD6 impairs chromatin remodeling at promoters via co-factor recruitment. (**a**) *In silico* rendering of the SANT-SLIDE domain structure of wild-type and mutant CHD6 on the basis of published Chd1 SANT-SLIDE structures. (**b**) NanoDSF melting profiles (first derivative of the 330/350 nm absorbance ratio; two replicates) of purified wild-type (*blue*) and I1600M (*green*) full-length CHD6 along a temperature gradient. *: significantly different mean; *P*<0.01, two-tailed unpaired Student’s t-test. (**c**) Overexpression of wild-type or mutant *CHD6* ORFs cloned into a piggybac vector (*top*) in iPSCs exemplified by Venus-GFP expression (*bottom*). (**d**) Network representation of GO terms associated with all proteins co-immunoprecipitating with wild-type CHD6 in untreated iPSCs. (**e**) Heatmaps showing enrichment or depletion of CHD6-interacting proteins associated with particular biological functions upon starvation, etoposide treatment or in untreated mutant and wild-type iPSCs. (**f**) Venn diagrams (*left*) showing the overlap between CHD6-bound genes and genes differentially-regulated upon *SMARCB1* knockdown in hESCs. Bar plots (*right*) showing significantly-enriched GO terms associated with CHD6-bound up- (*orange bars*) or down-regulated genes (*blue bars*). (**g**) Bar plots showing changes in SMARCC1 (BAF155) ChIP-qPCR enrichment (fold enrichment ±SD; *N*=2) from monoallelic (*grey*) or heterozygous mutant (*green*) compared to wild-type iPSCs; *: significantly different to wild-type; *P*<0.05, two-tailed unpaired Student’s t-test. (**h**) As in panel g, but for ACTL6A (BAF53).

Thus, we examined whether the HSS-mutation would result in folding that interferes with CHD6 protein-protein interactions. We generated stable doxycycline-inducible iPSCs overexpressing equal levels of HA-tagged wt and mut-CHD6 via random genomic integrations of each ORF using a piggybac vector (as in ref. 37; **Fig. 4c**). Following 24 h of induction, HA-CHD6 and its interacting partners were immunoprecipitated and subjected to quantitative label-free mass-spectrometry. wtCHD6 presented an interactome consisting of general and specialized transcription factors, other types of chromatin remodelers, cell cycle regulators, RNA-binding proteins related to RNA processing, and factors involved in rRNA biogenesis (**Fig. 4d**). We also generated CHD6 interactomes upon 2 h of starvation or 1 h of etoposide treatment, and monitored how mutCHD6 interactors compare to those of the wild-type protein. Strikingly, the interactomes of mutCHD6 displayed a strong and consistent loss of chromatin remodeling complex components, for example those belonging to the BAF/PBAF complexes, across replicates and conditions (**Figs 4e** and **S4j**). Importantly, this loss was not due to expression changes in these subunits as judged by our 3’-end-seq data.

Two types of canonical SWI/SNF complexes exist in humans: the BRG-/BRM-associated factor (BAF) and the Polybromo-associated BAF (PBAF), both of which are crucial for normal development and differentiation^46,47^. To address their connection to CHD6-mediated gene regulation, we reanalyzed transcriptome data from *SMARCB1(INI1)*-knockdown human embryonic stem cells^48^. 140 *SMARCB1*-regulated genes were also found bound by CHD6, >40% of which are downregulated and associate with lysosomal and vacuole organization and the p53 pathway. The rest are upregulated genes linked to RNA processing, protein stability, and chromatin modifiers (**Fig. 4f**). Similarly, reanalysis of hESC knockdown data for the SWI/SNF component BRG1^49^ identified 64 differentially-regulated genes that are also bound by CHD6. Remarkably, these are related to bone morphogenesis, odontogenesis, heart and head formation and thus relevant to HSS phenotypes (**Fig. S4k**). Finally, we directly tested some of these targets for recruitment of the ACTL6A (BAF53) and SMARCC1 (BAF155) subunits in both het- and mut-iPSCs/CMs by ChIP-qPCR, as interaction with both subunits was lost in mutCHD6 mass-spec data (**Fig. S4j**). We found that BAF/PBAF-subunit binding was indeed significantly diminished (**Fig. 4g,h**) in line with all aforementioned data. Together, this set of data suggest that the HSS mutation precludes CHD6 from recruiting additional necessary chromatin remodeling factors to its target loci. This can, in turn, prolong CHD6 residence at these sites and negatively affect transcriptional activation. Given that CHD6 mostly binds autophagy-related loci, the downstream impact is a dampened autophagy flux and an impaired DNA damage repair.

## DISCUSSION

Some CHD chromatin remodelers are well studied and causally linked to human pathologies^8,9,13,10^, but the roles of CHD6 remained for their most part elusive. To address this, we exploited a *de novo* mutation linked to the rare Hallermann-Streiff syndrome and modeled the disease using iPSCs. In summary, our work uncovers an essential housekeeping role for CHD6 in autophagy regulation, which explains its consistent expression across most human tissues (https://www.proteinatlas.org/ENSG00000124177-CHD6/tissue). In line with this role, CHD6 displays remarkable overlap of target loci across diverse cell types (iPSCs, NCCs, and CMs). A considerable fraction of these CHD6-bound loci is also TFEB targets, apparently already primed in resting cells for prompt transcriptional response to pro-autophagy stimuli.

The HSS-relevant I1600M mutation in CHD6 does not impact its ability to bind and remodel chromatin *in vitro*. Nonetheless, CHD6-bound genes in mutant cells fail to adequately respond to starvation or DNA damage stimuli. Moreover, *in vivo* overexpression of mutCHD6 results in obvious developmental malformations reminiscent of the HSS phenotype. We can now mechanistically attribute this to the inability of the mutant CHD6 SANT/SLIDE domain to interact with and recruit co-remodelers to its target promoters. In our proteomics data, the PBAF/BAF complexes are two major such co-players. The PBAF complex has already been shown to interact with CHD7 in order to direct NCC formation via enhancer regulation^46^. Interestingly, PBAF was recently implicated in stress responses^50^ and in euchromatic DNA lesion repair^51^. This aligns well the fact that CHD6 is exclusively found in euchromatin (see ref. 14 and our ChIP-seq data), but not at CHD7 sites. Thus, this constitutes a different regulatory leg. Moreover, reanalysis of *SMARCB1*- and *BRG1*-knockdown data from hESCs, both PBAF/BAF subunits operating on promoters and enhancers^47,52^, showed that they overlap CHD6 in regulating lysosomal, autophagy, and DNA damage response pathways. This highlights a requirement for multiple, competing or converging, remodeling activities acting at the same promoter to ensure its precise and timely regulation^53^.

In a recent report implicating it in the oxidative DNA damage response, CHD6 was shown to be stabilized via reduced degradation and to relocate rapidly to sites of DNA damage. This was mediated by its chromodomain and central core subregions, and by an N-terminal poly(ADP-ribose)-dependent motif. It also involved interactions with the oxidative stress response transcription factor NRF2^17^. Like Moore and co-workers, we saw no change in *CHD6* expression upon etoposide-induced DNA damage (or upon starvation). CHD6 did not redistribute to etoposide-induced DNA lesions, while only the CHD6 SANT/SLIDE domain and not its chromodomain and central core (ATPase/helicase) regions are mutated in our HSS model. Moreover, aberrant autophagy is a *bona fide* cancer hallmark^54^ and might affect how CHD6 functions in cancer cell lines. Thus, we suggest that the primary housekeeping role of CHD6 in normal human cells is autophagy control. It is also worth noting that a subset of CHD6-bound genes are involved in RNA metabolism and processing. CHD6 itself interacts with a number of RNA binding and processing factors in our proteomics data. Interactions with many of these factors are specifically hindered following etoposide treatment of CHD6-mutant cells. This aspect further links CHD6 to the transcriptional control of DNA damage responses and merits future investigation.

Finally, increased burden of un- or poorly resolved DNA damage is a hallmark of ageing, and mutations in the DNA damage response machinery give rise to premature ageing-like phenotypes, like in ERCC1^55^ or XPA-deficient cells^56^. In addition, mutations in particular SWI/SNF remodeling subunits critically affect stem cell functions and accelerate cellular ageing^57^. We observed acquisition of senescence-like features in HSS-mutant cells. CHD6 must operate upstream of these changes because it coordinates the autophagy and DNA damage responses across tissues in the developing organism. In support of this, the key regulator p62 normally mediates autophagy effects onto DNA repair mechanisms, likely in conjunction with p53^58^, but this mediation is gradually impaired with age and restored via lifespan-extending interventions^59^. In HSS-mutant cells, p62 levels are overall lower and the autophagy-to-DNA damage repair axis disturbed. Thus, we can now propose that HSS stems from autophagy-driven DNA damage burden and induction of a cellular ageing phenotype.

## METHODS

### Whole-exome sequencing (WES) and reprogramming of HSS fibroblasts

Exonic and adjacent intronic sequences were enriched from genomic DNA isolated from blood and salinva of the index patient and the parents using the NimbleGen SeqCap EZ Human Exome Library v2.0 enrichment kit, and sequenced on a HiSeq2000 platform (Illumina). WES data analysis and filtering of mapped targeted sequences was carried out using the “Varbank” exome and genome analysis pipeline v2.6 of the Cologne Center for Genomics (CCG, University of Cologne, Germany) and we obtained a mean coverage of 75-95 reads, with 95.9-96.7% of target sequences covered more than 10x. Trio-WES data was filtered for high quality (coverage of >6 reads, minimum quality score of 10), rare autosomal recessive and *de novo* variants (i.e., with minor allele frequency of <0.5% in the 1000 Genomes database, the Exome Aggregation Consortium browser, and not annotated in all in-house WES datasets of the CCG). Primary fibroblasts from the index patient were isolated via standard skin biopsy, and all downstream work was performed in accordance to the Helsinki Declaration protocols and reviewed and approved by the local institutional Ethics boards (University Hospital Cologne, Germany; University Medical Center Göttingen, Germany).

### iPSC culture and differentiation into cardiomyocytes or neural crest cells

iPSCs were cultured in FTDA media as previously described^60^. Once confluent, cells were dissociated using accutase at 37°C for 10 min (Sigma-Aldrich) and 450-600,000 cells were seeded per each well of a 6-well plate. Differentiation into cardiomyocytes was performed as previously described^61^. Briefly, confluent iPSCs were dissociated into single cells using accutase. Cells were then counted and 600,000 cells for each well of a 24-well plate were aliquoted and spun for 2 min at 300 x *g* at room temperature. The cell pellet was resuspended in ITS medium (knockout DMEM, 1x penicillin/streptomycin/glutamine, 1x ITS supplement, 10 μM Y-27632, 25 ng/ml FGF2, 1-2 ng/ml BMP4 and 1-2 μM CHIR99021) and seeded in matrigel-coated 24-well plates. Note that wt-iPSCs required 1.75 μM and 1.25 ng/ml, mut-iPSCs required 1.75 μM and 2 ng/ml, and patient-derived iPSCs required 1.75 μM and 1.75 ng/ml of CHIR and BMP4, respectively, for differentiation. To ensure the equal distribution and attachment of cells, plates were moved crosswise, tapped several times and left for 20 min at room temperature before being transferred into the CO_2_ incubator. After 24 h, ITS growth medium was replaced by TS medium [knockout DMEM, 1x penicillin/streptomycin/glutamine, 1x TS supplement (100x stock: 5.5 μg/ml Transferrin and 6.7 ng/ml Sodium Selenite in 100ml sterile PBS), and 250 μM 2-phospho-L-ascorbic acid]. After 48 h, TS medium was replenished and supplemented with 10 μM of the Wnt-inhibitor IWP-2 for 48 h. After 48 h, media were changed to fresh TS until beating cells were observed on day 8. At this point, medium is changed back to knockout DMEM supplemented with 2% FCS, L-glutamine and 1x penicillin/streptomycin for maintenance until cells are used for downstream analysis. Differentiation into neural crest cells was performed as previously described^62^. Briefly, iPSCs clusters were transferred into neural induction media [NIM; 1:1 ratio of DMEM-F12 (Invitrogen #10565-018) and Neurobasal media (Invitrogen #21103-049) complemented with 0.5x N2, 0.5x B27, 5 μg/ml insulin, 20 ng/ml βFGF, 20 ng/ml EGF, 1x pen/strep] in uncoated polypropylene dishes. Cell-spheres/-rosettes were allowed to spontaneously attach and after 6-9 days hNCLCs (human neural crest-like cells) migrated out of these. Subsequently, rosettes were dissected away and P0-isolated NCLCs were dissociated using accutase and re-plated for 10-15 days of maintenance in NIM in dishes coated with 1 μg/ml fibronectin.

### Generation of isogenic iPSC lines via CRISPR-Cas9 genome editing

gRNAs were designed against DNA stretches within exon 31 of the *CHD6* gene or the intron preceding it using an online tool (http://crispr.mit.edu). Each gRNA was assembled from two complementary oligonucleotides containing the “NGG” PAM sequence and distinct 4 bp-overhangs (“CACC” and “AAAC”) allowing for cloning into the *Bbs*I restriction of the pX330A vector as previously described^63^. Then, a single-stranded DNA oligo (ssODN)^64^ or a plasmid carrying homology arms^65^ were provided as templates for homologous recombination. iPSCs were seeded at low density in a 6-well plate (175,000 cells/well) and transfected 12 h later by the dropwise addition of a mixture of 200 μl OptiMEM (Invitrogen), 12 μl FuGene HD (Promega) and a total of 3 μg from all plasmids. After 24 h GFP expression was detectable microscopically; transfection efficiency was estimated to be ≥60%. Cells were then selected in 0.5 μg/ml puromycin for 24 h and reseeded in clonal dilution (5,000-8,000 cells/well of a 6-well plate). Individual clones were screened using PCR and, ultimately, insertion of the desired mutation was verified by Sanger sequencing. All gRNAs and ssODNs used in this study are listed in **Table S1**.

### CytoScan genome analysis of iPSC lines

Genome integrity of all genome-edited and reprogrammed iPSC lines was assessed using CytoScan HD microarray technology (ThermoFisher Scientific), which allows for the reliable detection of 25-50 kbp-long copy number changes genome-wide. For this analysis, intact genomic DNA was isolated using the Quick DNA Miniprep plus kit (Zymo Research) as per manufacturer’s instructions, and data analyzed via the ChAS suite v4.0 against the hg19 reference genome assembly. A summary is provided in **Table S2**.

### Generation and analyses of RNA-seq data

Cells from different genotypes/differentiations were harvested in Trizol (Life Technologies) and total RNA was isolated and DNase-treated using the Direct-zol RNA miniprep kit (Zymo Research) as per manufacturer’s instructions. In the case of cardiomyocytes, cells at day 10-11 of differentiation were used, while in the case of NCCs, cells were obtained by collecting migratory cells from dissected and re-plated rosettes. Barcoded cDNA libraries were generated using the TruSeq RNA library kit (Illumina) via selection on poly(dT) beads. The resulting libraries were paired-end sequenced to >50 million read pairs on a HiSeq4000 platform (Illumina). Transcript quantification was performed using Kallisto v0.44.0^66^, which pseudoaligns RNA-seq reads to transcripts (Ensembl annotation used: GRCh37, comprising all known protein-coding sequences from hg19). After transcript quantification, pseudo-counts were further processed via Sleuth^67^, which bootstraps Kallisto output to ascertain and correct for technical variation. To test for association between gene expression and genotype, we created a linear model and tested the effect of the *CHD6* mutation in each cell type via gene-level analysis. *P*-values were corrected for multiple hypothesis testing using the Benjamini-Hochberg algorithm and an FDR=0.05 as a significance threshold for all figures and tables. All data preprocessing and visualization was performed using R v3.3.3 and Bioconductor v3.4, and GO term enrichment analyses using Metascape (http://metascape.org/)^68^. Differentially-regulated genes per each line are listed in **Table S3**. For qPCR, the SYBR Green JumpStart Taq ReadyMix (Sigma-Aldrich) was used as per manufacturer’s instructions.

### 3’-end RNA sequencing and analysis

3’-end RNA-seq was used as a lower-cost alternative to mRNA-seq to interrogate multiple conditions and genotypes. Libraries were prepared from total RNA using the QuantSeq 3’ mRNA-Seq Library Prep Kit (Lexogen), and single-end sequenced on a HiSeq4000 platform (Illumina) generating ~15×10^6^ 100 nt-long reads per sample. Reads were quality assessed and mapped to hg19 using STAR^69^. Reads uniquely mapping to exons were quantified using HTSeq-count and differential gene expression was assessed using DESeq2^70^. Differentially-regulated genes per each cell line and treatment are listed in **Table S4**.

### ChIP-seq and data analysis

For each batch of ChIP experiments, ~12 million cells were crosslinked in 2% PFA for 45 min at 4°C. From this point onward, cells were processed via the ChIP-IT High Sensitivity kit (Active motif) as per manufacturer’s instructions, but using the NEXSON protocol for the isolation of nuclei^71^. Chromatin was sheared to 200-500-bp fragments on a Bioruptor Plus (Diagenode; 2x 20-26 cycles of 30s “on” and 30s “off” at the highest power setting), and immunoprecipitations were carried out by adding 4 μg of the appropriate antisera (CHD6, Bethyl A301-221A; TF3B, Bethyl A303-673A) to ~30 μg of chromatin and incubating on a rotator overnight at 4°C in the presence of protease inhibitors. Following addition of protein-A/G agarose beads and washing, DNA was purified using the ChIP DNA Clean & Concentrator kit (Zymo Research) and used in qPCR or sequencing on a HiSeq4000 platform (Illumina). qPCRs were performed with the primers listed in **Table S1**. Where ChIP-seq was performed, >20 million reads were obtained, also for the relevant “input” samples. Raw sequencing reads (typically 50 nt-long) were analyzed using the HiChIP pipeline^72^, and peaks were called using MACS2^73^. Thresholded CHD6 ChIP-seq peaks (q-value <0.05) per each cell type and genotype are listed in **Table S5**. For plotting ChIP-seq signal coverage over particular genomic regions, *ngs.plot* was used^74^.

### Immunostaining and imaging

Cells were grown on coverslips, fixed in 4% PFA/PBS for 10 min at room temperature, washed once in 1x PBS, permeabilized in 0.5% Triton-X/PBS for 5 min at room temperature, blocked in 1% BSA/PBS for 1 h before incubating with the primary antibody of choice for 2 h to overnight. Cells were next washed twice in 1x PBS for 5 min, before incubating with the appropriate secondary antisera for 1 h at room temperature. Nuclei were counterstained with DAPI (Sigma-Aldrich) for 5 min, washed, and coverslips mounted onto slides in Prolong Gold Antifade (Invitrogen). For image acquisition, a widefield Leica DMI 6000B with a HCX PL APO 63x/1.40 (Oil) objective was used, making sure exposure times were maintained constant across samples in each imaging session for the same immunostaining. Finally, images were analyzed using the Fiji suite^75^ as follows. First, background signal levels were subtracted using the embedded function (rolling ball function of 50-px radius with a sliding paraboloid and disabled smoothing), and the DAPI channel was used to determine the area of interest where signal would be quantified from. Measured mean signal intensities were used to generate plots in R or via Instant Clue^76^.

### Comet assays

Comet assays were performed as previously described^77^. Cells are treated with 30 μg/ml etoposide for 30 min, harvested with accutase to prepare a single-cell suspension, counted and diluted to a cell density of ~2 × 10^4^ cells/ml in PBS without bivalent cations on ice. Electrophoresis slides are covered with low-melting agarose, alkaline lysis is allowed to run overnight, before slides are submerged in rinse solution for 20 min, the solution exchanged another 2 times to ensure removal of salts and detergents, and finally submerged in an electrophoresis chamber filled with fresh wash buffer. Electrophoresis proceeds for 25 min at a constant current of 40 mA, before slides are neutralized in distilled water, placed in staining solution containing 2.5 μg/ml of propidium iodide for 20 min, and rinsed again in distilled water. For image analysis, *comet score* (http://rexhoover.com/index.php?id=cometscore) was used with doublets or comets at slide edges discarded from analysis. The length and intensity of DNA “tails” relative to “heads” are used as proxies for the amount of DNA damage in individual nuclei.

### Generation of stable CHD6 overexpression lines, immunoprecipitation, and proteomics

Full-length *CHD6* cDNA lacking the stop codon was PCR-amplified from wild-type NCC cDNA and cloned into a piggyback backbone (Ka0717_pPb-hCMV-cHA-IRES Venus)^78^ between the *Mlu*I and *Spe*I restriction sites, thus positioning the *CHD6* cDNA in frame with a C-terminal HA-tag. The HSS-relevant mutation (C4800G) was introduced to the wt*CHD6* piggybac vector using site-directed mutagenesis. Wild-type iPSCs were transfected as described above, using the wt- or mut*CHD6*-containing piggybac together with a transposase-expressing vector enabling few random integrations of the construct into the genome. Individual clones were selected after clonal dilution and selection on the basis of high Venus signal. *CHD6* overexpression was confirmed by immunofluorescence and western blots using anti-HA antisera. Following overexpression, wt/mutCHD6 immunoprecipitation was performed on freshly-harvested doxycycline-induced iPSCs. First, cell nuclei were isolated by incubating cells for 15 min on ice in NIB buffer (15mM Tris-HCL pH 7.5, 60 mM KCl, 15 mM NaCl, 5 mM MgCl_2_, 1mM CaCl_2_, 250 mM sucrose) containing 0.3% NP-40. Nuclei were pelleted for 5 min 800 x *g* at 4°C, washed twice in the same buffer, lysed for 10 min on ice in IP buffer (150 mM LiCl, 50 mM Tris-HCl pH 7.5, 1mM EDTA, 0.5% Empigen) freshly supplemented with 2 mM sodium vanadate, 1x protease inhibitor cocktail (Roche), PMSF (10 μl), 0,5mM DTT, 5 μl Caspase inhibitor III (Calbiochem), and 50 units Benzonase per ml of IP buffer, before preclearing cell debris by centrifugation at >15,000 x *g* at 4°C. Finally, 1 mg of the lysate was incubated with anti-HA antisera overnight at 4°C. Magnetic beads (Active Motif) were then washed once with 1x PBS-Tween and combined with the antibody-lysate mixture. Following a 2-h incubation at 4°C, beads were separated on a magnetic rack and washed 5x, 5 min each in wash buffer (150 mM KCl, 5 mM MgCl_2_, 50 mM Tris-HCl pH 7.5, 0.5% NP-40) and another two times in wash buffer without NP-40. Captured proteins were predigested and eluted from the beads using digestion buffer (2M Urea, 50 mM Tris-HCl pH 7.5, 1 mM DTT) supplemented with trypsin and eluted from the beads with elution buffer (2M Urea, 50 mM Tris-HCl pH 7.5, 5 mM chloroacetamide) supplemented with trypsin and LysC, before subjected to mass-spectrometry on a Q-Exactive platform (ThermoFisher). The full list of peptide hits and their analysis is provided in **Table S6**.

### Western blotting

Western blotting was performed as previously described^37^. In brief, ~2×10^6^ cells were enzymatically lifted or gently scraped off cell culture dishes, and pelleted for 5 min at 600 x *g*. The supernatant was discarded, and the pellet resuspended in 100 μl of ice-cold RIPA lysis buffer containing 1x protease inhibitor cocktail (Roche), incubated for 30 min on ice, and centrifuged for 15 min at >15,000 x *g* to pellet cell debris and collect supernatant. Total protein concentrations were determined using the Pierce BCA Protein Assay Kit (ThermoFisher Scientific), before extracts were stored at −80°C. Proteins were resolved by SDS-PAGE, transferred onto membranes using the TransBlot Turbo setup (Bio-Rad), and detected using the antisera and dilutions listed in **Table S7**.

### MNase nucleosome isolation and qPCR

iPSCs were grown in 6-well plates to ~80% confluency and rinsed with PBS prior to addition of 1 ml of freshly prepared permeabilization buffer (15mM Tris/HCl pH 7.6; 60 mM KCl; 15 mM NaCl; 4 mM CaCl2; 0.5 mM EGTA; 300 mM sucrose; 0.2% NP-40; 0.5 mM b-mercaptoethanol supplemented with MNase (company) at 1u/ml final concentration. MNase was added for 3 min at 37°C, and stopped by addition of an equal volume of stop buffer (50 mM Tri/HCl pH 8.0; 20 mM EDTA; 1% SDS). Finally, 250 μg RNase A were added for 2 h at 37°C, followed by addition of 250 μg proteinase K and incubation at 37°C overnight. Next day, DNA was isolated via standard phenol-chloroform extraction, digestion efficiency was determined after electrophoresis in 1% agarose gels, and mononucleosomes were isolated from the gel. Nucleosome occupancy was assessed with qPCR using the primers listed in **Table S1**.

### β-galactosidase staining, acidic organelle detection with LysoTracker, and FACS analysis

Senescence-associated β-galactosidase assays (Cell Signaling) were performed as per manufacturer’s instructions before manually counting positive cells Acidic organelles were detected using Lysotracker Deep Red (ThermoFisher) as per manufacturer’s instructions. Cell cycle analysis was performed using live cell nuclear staining with RedDot1 (Biotum) as per manufacturer’s recommendations. Flow cytometry data were analyzed via the FlowJo software (https://www.flowjo.com/).

### In vitro and in vivo migration assays

For *in vitro* migration (scratch) assays, wild-type and patient-derived NCCs were grown in 6-well plates and one scratch per well was manually inflicted using a sterile cell scraper. Cell migration into the scratch was monitored for up to 8 h in 2-h increments by brightfield microscopy. For *in vivo* migration assays, GFP-labeled wild-type and mutant iPSCs were differentiated into NCCs as described above. Using a blunt glass capillary, NCCs were lifted with the help of accutase and inserted into the developing anterior neural region (i.e., midbrain) of chicken embryos at stage HH10. Operated eggs were resealed with medical tape and incubated until stages HH20, when embryos were isolated and fixed in 4 % PFA for immunofluorescence; GFP was detected using anti-GFP antisera (TP401, OriGene).

### In vivo CHD6 overexpression in chicken embryos

For *in vivo* CHD6 overexpression, electroporation of chicken embryos was performed as previously described^79^. Briefly, eggs of stage HH9-10 were windowed, and the extraembryonic membrane partially removed. A vector containing mutant or wild type *CHD6* ORFs (Ka0717_pPb-hCMV-cHA-IRES Venus-CHD6m and Ka0717_pPb-hCMV-cHA-IRES Venus-CHD6m; *this paper*) were first mixed 9:1 with the plasmid encoding the transposition transactivator, and this mixture then mixed 2:1 with Fast Green solution (Sigma) and microinjected into the neural tubes of the embryos. The neural tube was then electroporated with 5x square pulses of 20 V/20 ms using the Intracel TSS20 Ovodyne electroporator, eggs were resealed with tape and re-incubated for 24 h, before tape was removed and 10 μl of 2 μg/ml doxycycline were added at the site of electroporation and eggs were sealed again and re-incubated until stage HH20. CHD6 overexpression was confirmed in transgenic embryos on the basis of GFP signal under an Olympus SZX16 stereomicroscope with an EXFO X-cite series 120PC Q apparatus.

### NanoDSF CHD6 thermal stability determination

For thermal unfolding experiments, full-length wildtype and I1600M CHD6 protein preparations were diluted to a final concentration of 150 nM in EX80 buffer, and 10 μL of each sample per capillary was prepared. The samples were loaded into high sensitivity UV capillaries, and measurements were carried out on a Prometheus NT.48 instrument with a temperature gradient set to increase 1°C/min in the range of 20°C to 90°C and independently replicated twice.

### CHD6 protein expression and purification and in vitro assays

Steps from cloning to protein expression and purification were carried out as described previously^80^. After introducing a C-terminal FLAG-tag to the CHD6 cDNA, the CHD6-I1600M mutation was generated using the QuikChange Site-Directed Mutagenesis kit (ThermoFisher) according to the manufacturer’s instructions. Full-length wt and mutant clones were used as templates to generate SANT-SLIDE 2 domain constructs by PCR. All baculoviruses were produced according to the Bac-to-Bac Baculovirus Expression Systems manual (Life Technologies) and CHD6-FLAG protein constructs were next purified from ~4 L of Sf9 insect cell cultures. Purified proteins were used in electrophoretic mobility shift assays (EMSA), as well as in nucleosome mobilization/sliding and restriction enzyme accessibility (REA) assays using DNA and nucleosome substrates as described previously^80^. Detection and quantification of non-radioactive substrates was achieved following staining with SYBR Gold Nucleic Acid Gel Stain (ThermoFisher).

### Statistical analyses

*P*-values associated with Student’s t-tests were calculated using GraphPad (http://graphpad.com/) and R, and those associated with the Wilcoxon-Mann-Whitney test using the EDISON-WMW online tool (https://ccb-compute2.cs.uni-saarland.de/wtest/). Unless otherwise stated, only *P*-values <0.01 were deemed as significant.

### Data availability

All sequencing data generated in this study have been deposited in the NCBI Gene Expression Omnibus (GEO) repository under the accession numbers GSE135832 and GSE136057.

## Author contributions

YK, AZ, JB, JT, GL and KB performed experiments; SF, NR and LK assisted with cardiomyocyte differentiations and CRISPR/Cas9 editing; RR, ML and ARI generated *in vivo* data and assisted with neural crest cell differentiations; YK, EGG and NJ undertook all bioinformatics analyses; JA and PN performed high throughput sequencing; HK, YL, GY, AH, FJK, EW, HB, and BW provided patient samples and WES data; YK and AP conceived the study; YK, KB, and AP wrote the manuscript with input from all co-authors.

## Acknowledgements

We would like to thank all members of the Papantonis, Kurian, Rada-Iglesias, and Wollnik laboratories for helpful discussions, Ioanna Papadionysiou for help in ChIP assays, and Alexander Brehm for critical reading of the manuscript. We would also like to thank the CMMC Imaging and FACS sorting facilities, the CECAD Proteomics facility, the Cologne Center for Genomics, and the High-Performance Computing cluster “CHEOPS” of the University of Cologne for continuous access to resources. This work was supported by UKGM (Project-Nr. 5/2016) and the Deutsche Forschungsgemeinschaft via TRR81/3, as well as by CMMC core funding and an Else-Kroener-Fresenius-Stiftung “Key-Project” grant (2015_A125).

