## Supplemental Figs S1-S4 for "Overarching control of autophagy and DNA damage response by CHD6 revealed by modeling a rare human pathology"

### CONTENTS

—This Supplement includes **Figures S1-S4** and legends for **Tables S1-S7** (that are provided as *.xlsx* files)

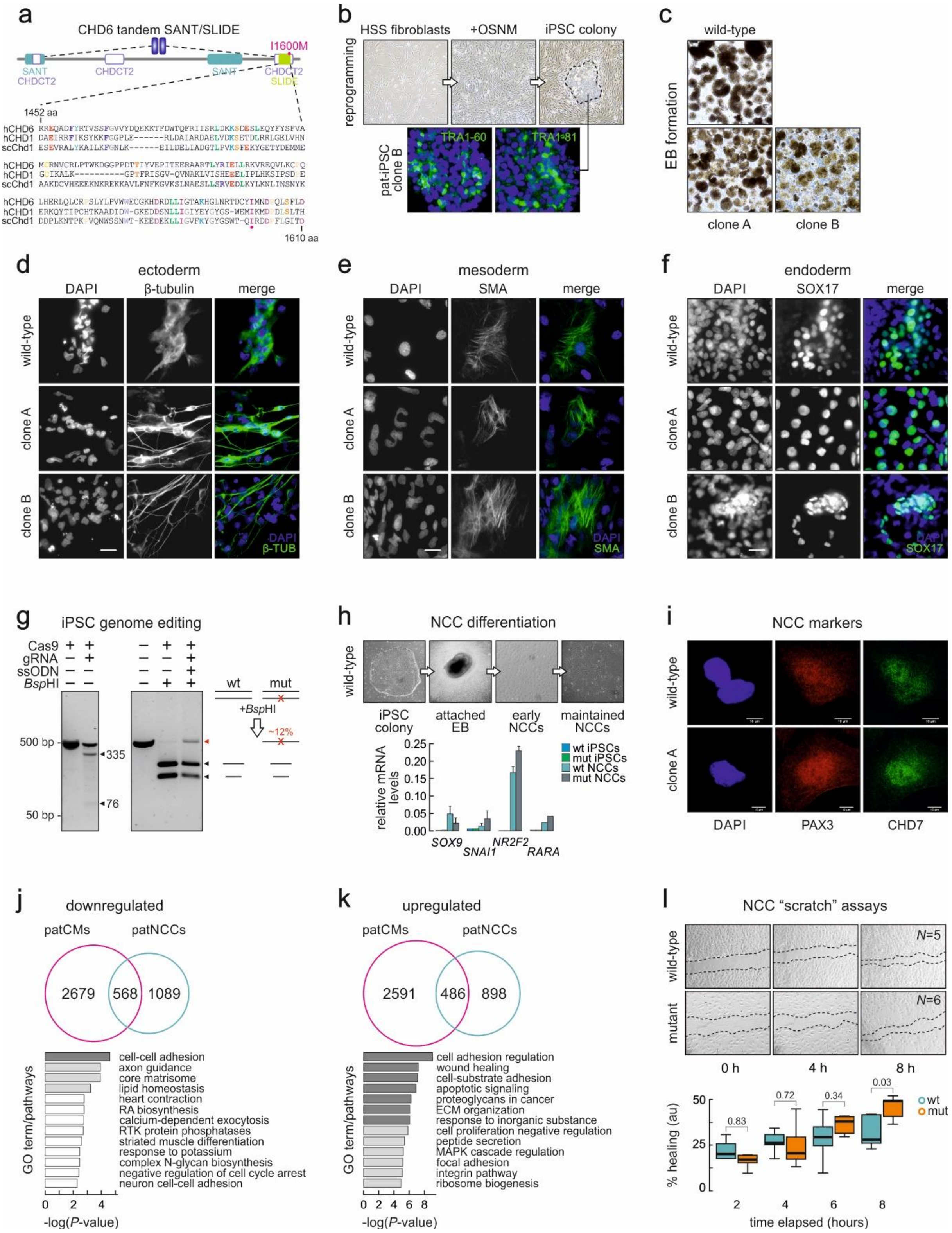

**Fig. S1. Reprogramming, genome editing, and differentiation of iPSC lines.**

(**a**) Identification of domains within the tandem SANT/SLIDE domain of CHD6 by alignment to resolved human (h) and yeast (*Sacharomyces cerevisiae*, sc) CHD1 domains in the PDB database.

(**b**) Representative brightfield images of patient-derived fibroblasts (*top left*) undergoing reprogramming via OCT4, SOX2, NANOG, and MYC overexpression (*top middle*), before single iPSC colonies are picked from the plate (dashed line; *top right*). Representative immunofluorescence

images of patient-derived iPSC clones stained for the TRA1-60 (*bottom left*) and TRA1-81 markers (*bottom right*) in nuclei are counterstained with DAPI. Bar: 10 µm.
(**c**) Representative brightfield images of wild-type (*top*) and patient-derived iPSCs (*bottom*) giving rise to embryonic bodies (EBs).
(**d**) Representative immunofluorescence images of wild-type and two patient-derived iPSC clones (*middle* and *bottom rows*) efficiently differentiated into ectoderm and stained for β-tubulin expression. Bar: 10 µm.
(**e**) As in panel d, but differentiated into mesoderm and stained for SMA expression. Bar: 10 µm.
(**f**) As in panel d, but differentiated into endoderm and stained for SOX17 expression. Bar: 10 µm.
(**g**) Agarose gel electrophoresis profiles of iPSCs cut by Cas9 at the CHD6 locus (*left*) and of a wild-type and an edited iPSC population diagnosed for carrying the HSS mutation via *Bsp*HI restriction digest.
(**h**) Representative brightfield images of iPSCs differentiated into NCCs (*top*) and RT-qPCR data of NCC marker genes (mean ±S.D., n=3; *bottom*).
(**i**) Representative immunofluorescence images of wild-type and mutant iPSC-derived NCCs stained for the PAX6 (*left*) and CHD7 markers (*right*). Nuclei are counterstained with DAPI. Bar: 10 µm.
(**j**) Bar plot showing the top significantly enriched GO terms/pathways associated with downregulated genes (at least 0.6 $log_2$-fold change, $P_{adj}$<0.05) shared by patient-derived CMs and NCCs.
(**k**) As in panel j, but for upregulated genes shared by patient-derived CMs and NCCs.
(**l**) *In vitro* migration (“scratch”) assays of wild-type and mutant NCCs and their quantification (*below*) over a time course of 8 h; two-tailed Student’s t-test *P*-values for each time point are also shown.

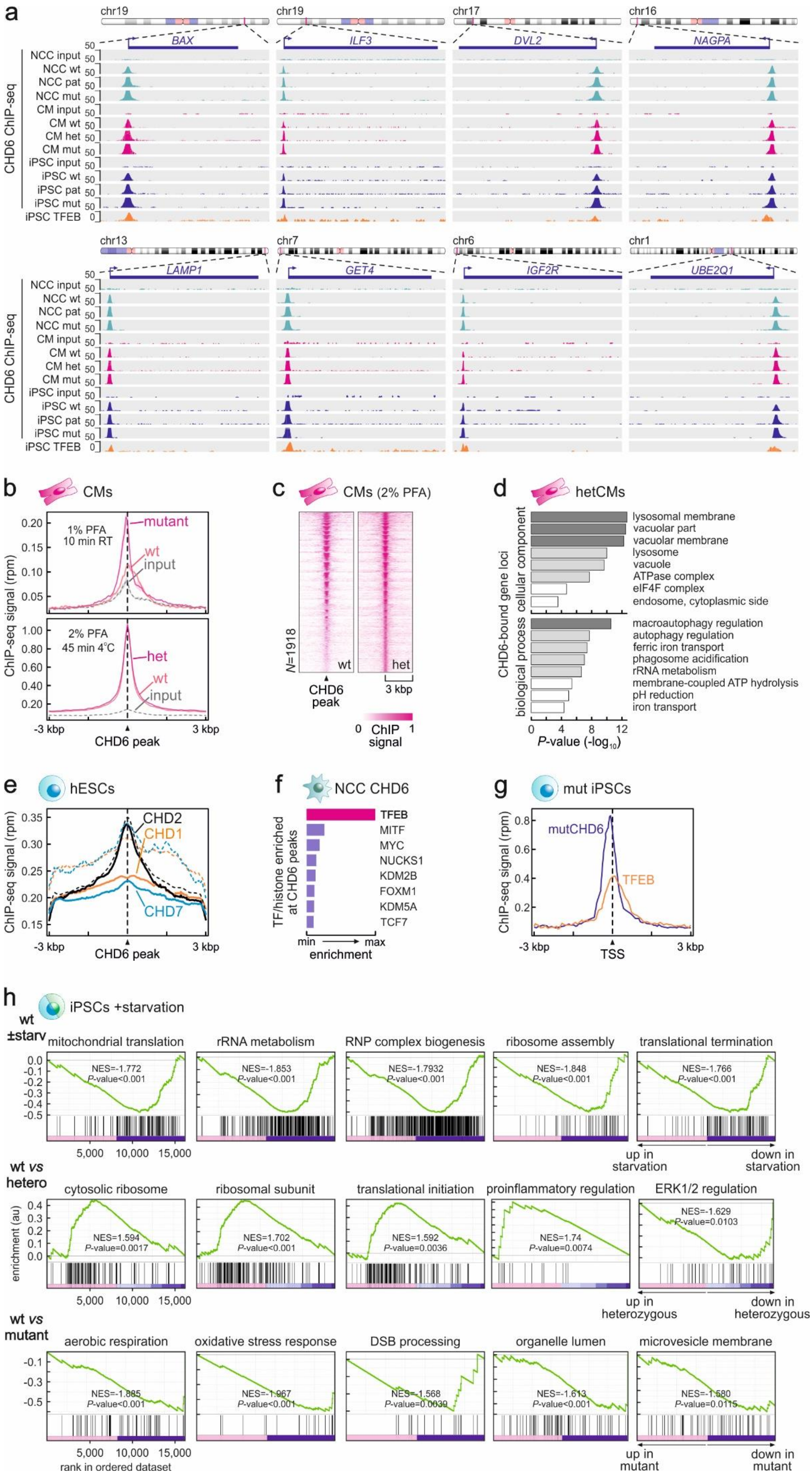

a
CHD6 ChIP-seq
NCC input
NCC wt
NCC pat
NCC mut
CM input
CM wt
CM het
CM mut
iPSC input
iPSC wt
iPSC pat
iPSC mut
iPSC TFEB
BAX
ILF3
DVL2
NAGPA
LAMP1
GET4
IGF2R
UBE2Q1
b
CMs
1% PFA 10 min RT
2% PFA 45 min 4°C
mutant
wt
input
het
ChIP-seq signal (rpm)
CHD6 peak
c
CMs (2% PFA)
N=1918
wt
het
3 kbp
ChIP signal
d
hetCMs
CHD6-bound gene loci
cellular component
biological process
lysosomal membrane
vacuolar part
vacuolar membrane
lysosome
vacuole
ATPase complex
eIF4F complex
endosome, cytoplasmic side
macroautophagy regulation
autophagy regulation
ferric iron transport
phagosome acidification
rRNA metabolism
membrane-coupled ATP hydrolysis
pH reduction
iron transport
P-value (-log10)
e
hESCs
CHD2
CHD1
CHD7
f
NCC CHD6
TF/histone enriched at CHD6 peaks
TFEB
MITF
MYC
NUCKS1
KDM2B
FOXM1
KDM5A
TCF7
min
max
enrichment
g
mut iPSCs
mutCHD6
TFEB
TSS
h
iPSCs +starvation
wt ±starv
mitochondrial translation
rRNA metabolism
RNP complex biogenesis
ribosome assembly
translational termination
NES=-1.772 P-value<0.001
NES=-1.853 P-value<0.001
NES=-1.7932 P-value<0.001
NES=-1.848 P-value<0.001
NES=-1.766 P-value<0.001
up in starvation
down in starvation
wt vs hetero
cytosolic ribosome
ribosomal subunit
translational initiation
proinflammatory regulation
ERK1/2 regulation
NES=1.594 P-value=0.0017
NES=1.702 P-value<0.001
NES=1.592 P-value=0.0036
NES=1.74 P-value=0.0074
NES=-1.629 P-value=0.0103
enrichment (au)
up in heterozygous
down in heterozygous
wt vs mutant
aerobic respiration
oxidative stress response
DSB processing
organelle lumen
microvesicle membrane
NES=-1.885 P-value<0.001
NES=-1.967 P-value<0.001
NES=-1.568 P-value=0.0039
NES=-1.613 P-value<0.001
NES=-1.580 P-value=0.0115
rank in ordered dataset
up in mutant
down in mutant

**Fig. S2. Analysis of CHD6 ChIP-seq data from multiple cell types.**

(**a**) Representative genome browser views of CHD6 ChIP-seq in wild-type (wt), monoallelic mutant (mut) or patient-derived (pat) NCCs (*green*), CMs (*magenta*), and iPSCs (*blue*) aligned to iPSC-derived TFEB ChIP-seq data.

(**b**) Line plots showing ChIP-seq signal in the 6 kbp around CHD6-bound sites from wild-type (*light pink*) and homozygous-/heterozygous-mutant CMs (*magenta*); input signals serve as control (*grey*). ChIP in the top panel was performed using 1% PFA for 10 min at room temperature, compared to 2% PFA for 45 min at 4°C used in the bottom panel (yielding 271 peaks vs 642 wt-peaks, respectively).

(**c**) Heatmaps showing ChIP-seq signal distribution in the 6 kbp around CHD6-bound sites from wild-type (wt) or heterozygous mutant CMs (het) for all 1918 peaks detected.

(**d**) Bar plots showing the top significantly-enriched GO terms/pathways for CHD6-bound genes in heterozygous mutant CMs.

(**e**) Line plot showing CHD1 (*orange*), CHD2 (*black*), and CHD7 (*blue*) ChIP-seq signal in the 6 kbp around CHD6-bound sites from mutant iPSCs; signal distribution from the peak list of each dataset serves as positive control (*dashed lines*).

(**f**) Bar plots showing relative enrichment of signal from ENCODE transcription factors ChIP-seq at CHD6-bound sites from mutant NCCs.

(**g**) Line plots showing mutCHD6 (*blue*) and TFEB ChIP-seq signal (*orange*) in the 6 kbp around co-bound TSSs from mutant iPSCs.

(**h**) Gene set enrichment analysis of ranked gene expression data from 3'-end RNA-seq data of serum-starved wild-type compared to serum-starved heterozygous (*middle row*) or homozygous CHD6-mutant iPSCs (*bottom row*). The same analysis on wild-type iPSCs in the presence and absence of starvation provides a control (*top row*).

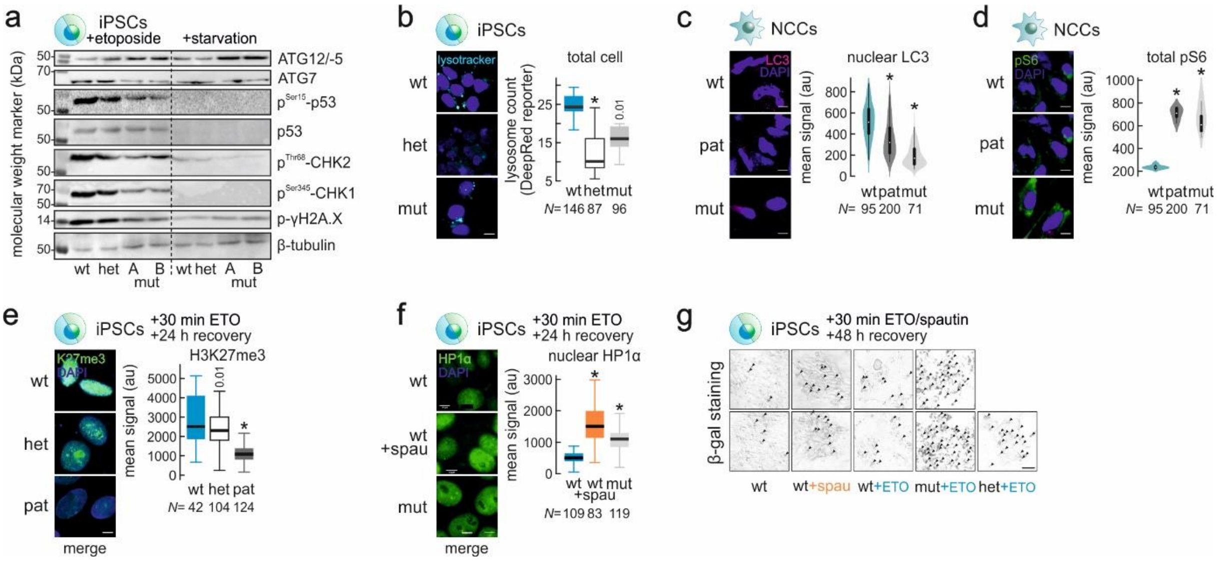


**Fig. S3. Autophagy, DNA damage, and senescence responses in *CHD6*-mutant cells.**

(**a**) Western blot analysis of proteins involved in autophagy and the DNA damage response in isogenic wild-type (wt) or two monoallelic-mutant iPSC clones (mutA,B) following etoposide treatment (*left*) or starvation (*right*).

(**b**) Representative immunofluorescence images (*left*) of wild-type (wt), monoallelic- (mut), or heterozygous-mutant (het) iPSCs stained for lysosomes (*light blue*) using an *in vivo* Lysotracker DeepRed reporter, and bean plots quantifying signal (*right*). The number of cells analyzed (*N*) is given below each plot. *: significantly different to wild-type; $P<0.01$, Wilcoxon-Mann-Whitney test.

(**c**) As in panel b, but for nuclear LC3 signal in NCCs.

(**d**) As in panel b, but for total phospho-S6 signal in NCCs.

(**e**) As in panel b, but for H3K27me3 signal in iPSCs treated for 30 min with etoposide and allowed to recover for 24 h.

(**f**) As in panel e, but for HP1α signal in iPSCs.

(**g**) Staining for β-galactosidase activity in wild-type, homozygous, and heterozygous mutant iPSCs treated or not with etoposide and/or spautin and allowed to recover for 48 h; β-gal positive cells are indicated (*arrows*).

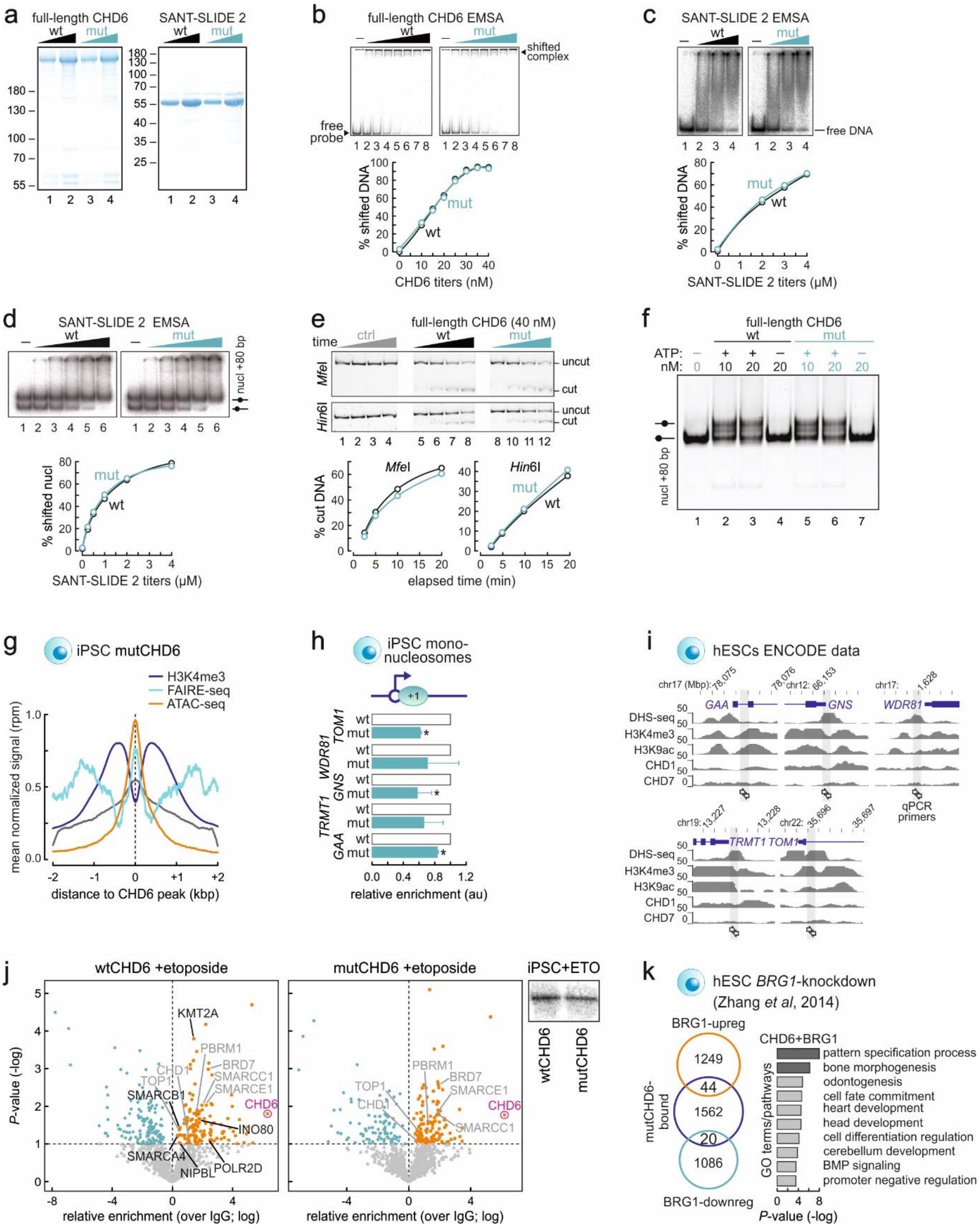

**Fig. S4. Effects of the I1600M *CHD6* mutation on gene expression and chromatin binding.**

(**a**) Coomassie-stained SDS-PAGE profiles of purified wild-type (wt) and I1600M (mut) full-length CHD6 and its SANT-SLIDE domain. The migration profiles of molecular weight markers (in kDa) are indicated.

(**b**) EMSAs performed with increasing titers of wild-type (wt) or mutant full-length CHD6 domain (mut) on a DNA template.

(**c**) As in panel b, but for the wild-type (wt) or mutant SANT-SLIDE 2 CHD6 domain (mut).

(**d**) As in panel c, but with increasing titers of the wild-type (wt) or mutant SANT/SLIDE 2 CHD6 domain (mut) on a nucleosomal template.

(**e**) Restriction enzyme accessibility assays (REA; *top*) and quantification (*bottom*) on a mononucleosome with time in the presence of 40 nm of wild-type (wt) or mutant full-length CHD6 (mut).
(**f**) Gel image of sliding assays using an end-positioned nucleosome and increasing titers of wild-type (wt) or mutant full-length CHD6 (mut).
(**g**) Line plot showing mean distribution of H3K4me3 ChIP-seq (*blue*), FAIRE-seq (*green*), and ATAC-seq signal (*orange*) in the 4 kbp around mutCHD6 binding sites in iPSCs.
(**h**) Bar plots showing mean relative enrichment of MNase-qPCR signal (±SD) at exemplary CHD6-bound genes using mononucleosomal DNA from iPSCs. *: significantly different to wild-type levels; $P<0.01$, two-tailed unpaired Student's t-test ($N=4$).
(**i**) Genome browser views showing hESC ENCODE DHS- and ChIP-seq data around five CHD6-bound gene promoters. The positions of primers used in MNase-qPCR (Fig. S4H) are indicated by arrows.
(**j**) Volcano plots of proteomics data following co-immunoprecipitation of wild-type (*left*) or mutant CHD6-interacting proteins (*middle*) obtained in iPSCs after etoposide treatment for 2 h. Proteins enriched significantly compared to non-specific IgG controls are shown in orange, and chromatin remodeling subunits lost in the mutCHD6 interactome are highlighted (*black*) over shared interactors (*light grey*); enrichment of CHD6 in the data (*magenta*) serves as positive control. Levels of wild-type and mutant CHD6 from iPSCs treated with etoposide are also shown (*right*).
(**k**) Venn diagrams (*left*) showing the overlap between CHD6-bound genes and genes differentially-regulated upon *BRG1* knockdown in hESCs. Bar plots (*right*) showing significantly-enriched GO terms associated with CHD6-bound up-/down-regulated genes.

### SUPPLEMENTAL TABLES

**Table S1.** Sequences of oligonucleotides used as gRNAs, repair templates, and primers in PCR/qPCR reactions (provided as an *.xlsx* file).

**Table S2.** Summary of microarray-based karyotyping data of all iPSC lines used in this work (provided as an *.xlsx* file).

**Table S3.** Genes significantly up- or downregulated in total RNA-seq data from patient-derived or homozygous mutant CMs and NCCs compared to their wild-type counter parts (provided as an .xlsx file).

**Table S4.** Genes significantly up- or downregulated in 3’ end RNA-seq data generated from wild-type, patient-derived or homozygous mutant iPSCs after serum starvation or etoposide treatment (provided as an *.xlsx* file).

**Table S5.** Lists of significant CHD6 ChIP-seq peaks and their coordinates (hg19) from all the different genetic backgrounds and cell types (provided as an *.xlsx* file).

**Table S6.** List of peptide hits and statistical analysis of proteomics data from wild-type and mutant iPSCs untreated, treated with etoposide or subjected to starvation (provided as an *.xlsx* file).

**Table S7.** Full list of antisera used and their working dilutions (provided as an *.xlsx* file).
